# Molecular Basis of β2 Integrin Activation by Talin Unveils Species-Specific Mechanisms of Integrin Signaling

**DOI:** 10.1101/2024.05.28.596271

**Authors:** Tong Gao, Nicholas A. Maskalenko, Salvin Kabir, Kerry S. Campbell, Jinhua Wu

**Affiliations:** Cancer Signaling and Microenvironment Program, Fox Chase Cancer Center, Philadelphia, PA 19111, USA; Department of Biology, College of Science & Technology, Temple University, PA 19122, USA

## Abstract

Integrins consist of 24 species, each with unique tissue-expression profiles and distinct biological functions. The β subunit of integrin interacts with the FERM-folded head domain of talin through an N-P-x-Y/F motif, triggering integrin activation. Although this motif is conserved across most integrin-β subunits, the precise molecular mechanisms governing talin’s selective recognition of different integrin species remains unclear. We determined the crystal structure of talin head in complex with the β2-integrin tail. The structure reveals a two-mode configuration featuring a “rocking” motion of the talin head FERM domain compared with its interaction with β3 integrin, resulting in distinct inter-subdomain interactions and unique cavities. Switching of the talin:β2 binding mode to the talin:β3 binding mode enhances β2-integrin affinity and boosts LFA-1-mediated natural killer cell cytotoxicity. Moreover, stabilizing of the C-terminal α-helix in the talin head enhances its affinity to integrin and its activation. Together, our data elucidate the structural basis by which talin orchestrates its function in mediating integrin activation in a species-specific manner.

**Significance statement:** Talin exhibits significantly lower affinity with lymphocyte-rich β2 integrins compared with β3 integrins. Our results unveil the configurational preferences of the talin head when engaged β2 and β3 integrins. We introduce a two-mode seesaw model wherein the talin head adapts specific binding modes in response to distinct integrin species. The two configurations differ in inter-subdomain interactions, revealing unique cavities and distinct binding dynamics in each binding mode. Thus, our findings present exciting opportunities of the development of species-specific therapeutic agents targeting integrin activity more precisely by orchestrating the structural dynamics of talin.

## Introduction

Integrins are a family of heterodimeric adhesion receptors that play a crucial role in cell adhesion and signaling. Human possesses 24 integrin species, formed by eighteen types of α-subunits and eight types of β-subunits^1^. These receptors are expressed in a tissue-specific manner and serve distinct biological functions upon activation and clustering of integrin^1,2^. This process is facilitated by talin and its cofactor, kindlin, through interacting with the cytoplasmic segment of integrin β-subunit^3^. Activated integrins serve functions essential for cell adhesion, migration, and organ development^2,4^. Integrin activity is further orchestrated, in a species-specific manner, by posttranslational modification and isoform diversity of the activators^5–7^, as well as other intracellular regulators such as migfilin, filamin, and paxillin^8–10^. Considerable efforts have been dedicated to exploring the molecular mechanisms underlying how various species of integrins are regulated and coordinated to precisely serve their specific functions^1,2,11–13^. Thus, the insights into the species-specific activation of integrin are of great significance for developing therapeutic agents that target integrin with greater precision.

All integrin β subunits except β4 and β8 integrins possess a highly conserved talin-binding site with an N-P-x-Y motif and a loosely conserved kindlin-binding site with the same motif (**Fig. 1A**). While talin is better known for its function of activating integrins via the “inside-out” pathway, it also connects integrins to cytoskeletal proteins such F-actin and vinculin, promoting integrin outside-in signaling that regulates focal adhesion assembly, cell differentiation, proliferation, and survival^14^. Integrins position this motif and preceding residues in an integrin-binding groove in the talin head domain (THD) to achieve strong affinity^15–17^. Among all β-subunits, β1, β2, and β3 integrins have been extensively studied due to their diverse functions across various cell types and their significant roles in disease pathologies. β1 is expressed ubiquitously and engages in various functions when partnering with different α-subunit partners. β2 integrin is exclusively expressed on leukocytes, playing crucial roles in regulating immune responses^18^. Specifically, αLβ2 (lymphocyte function-associated antigen-1, or LFA-1) is essential for lymphocyte trafficking, adhesion to endothelial cells, and formation of immunological synapses with targets of cytotoxic lymphocytes; αMβ2 (macrophage-1 antigen or Mac-1) mediates macrophage adhesion during inflammation^19,20^. β3 is predominantly found in platelets and endothelial cells. Integrin αIIbβ3 is a platelet-specific integrin curial for platelet aggregation and adhesion to ligands around vascular injury, thus facilitating thrombus formation and hemostasis ^4,21^, whereas integrin αvβ3 is pivotal in diverse processes including angiogenesis, bone resorption, and neovascularization^22^. However, antagonist inhibitors of integrin αvβ3 exhibit high toxicity and poor efficacy, likely due to the unwanted agonism, ligand-free signaling, and impaired functions in inducing apoptosis and angiogenesis^23^.

**Figure 1.**
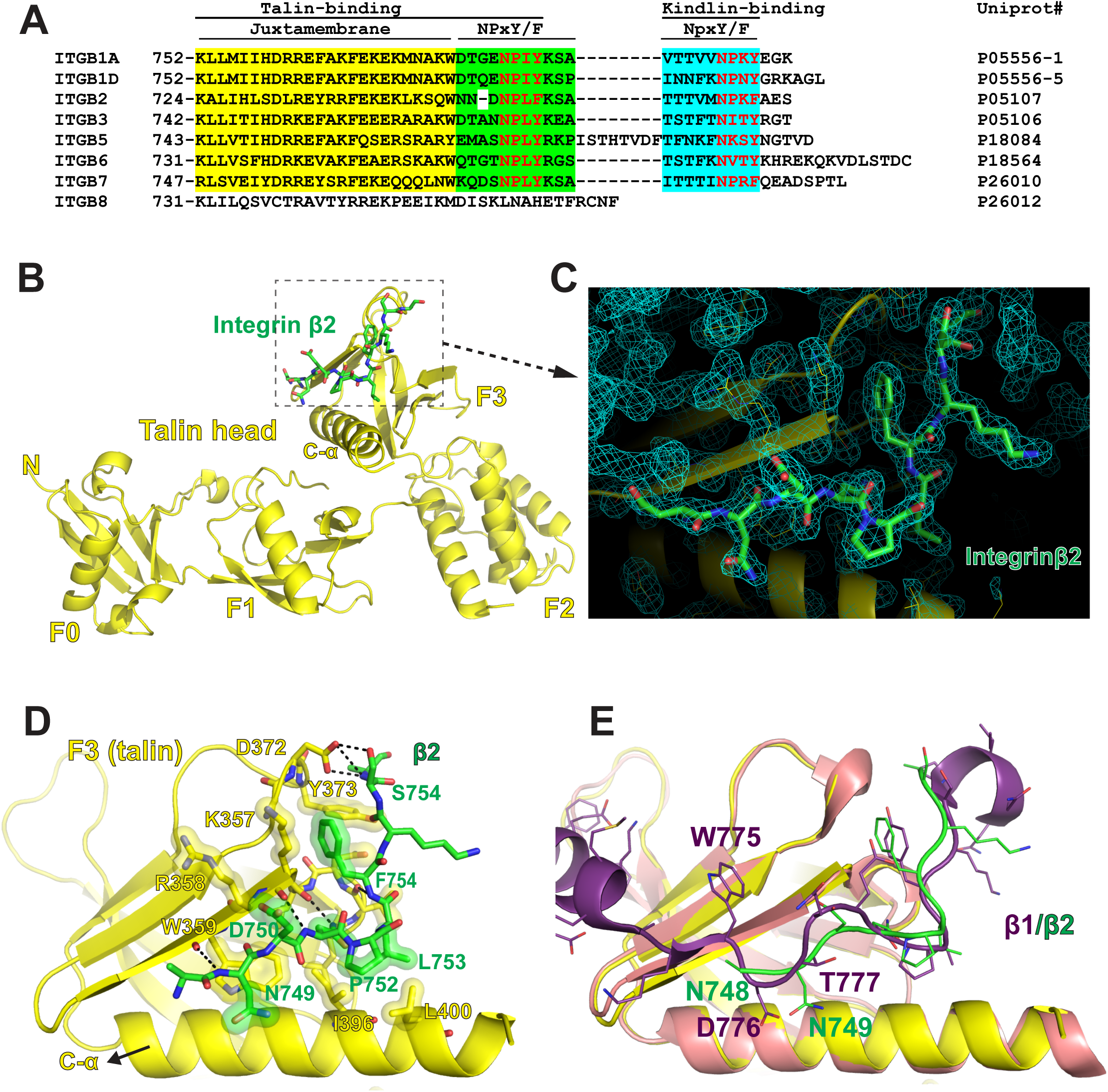
Crystal Structure of talin head domain in complex with integrin β2 NPxY/F motif. **A.** Sequence alignment of the cytoplasmic tails of integrin β subunits. The Juxtamembrane is highlighted in yellow, the NpxY/F motifs, colored in red, in the talin-binding site and the kindlin-binding site are highlighted in green and cyan, respectively. Uniprot codes of each integrin β subunit in human are shown. **B.** Structure of talin head domain in complex with integrin β2 is shown in cartoon presentation. **C**. Electron density of the NPxY/F motif in integrin β2 is contoured at σ=1.5. **D**. Side-chain interactions between talin-F3 and integrin β2. Ionic interactions are indicated by dotted line. Other interactions are indicated by surface representation. C-α helix is shown by the arrow. **E.** Superposition of the integrin β2-bound talin-F3 and the β1-bound talin-F3 with β1 in purple, β1-bound talin-F3 in salmon, β2 in green, and β2-bound talin-F3 in yellow, respectively.

Most integrin β subunits possess a cytoplasmic tail containing one talin-binding site and one kindlin-binding site, each harboring an N-P-x-Y/F motif. Crystal structure of talin head in complex with integrin β3 reveals a classic FERM-domain conformation^24^. A C-terminal poly-lysine motif, conserved in all FERM-domain proteins, stabilizes the FERM domain by mediating inter-subdomain interactions and secures a strong association with the cytoplasmic segment of integrin β3^24^. Moreover, the structure of integrin β1 in complex with a truncated talin head containing the F2-F3 subdomains in complex with integrin β1 reveals a similar integrin-binding groove in the F3 subdomain^16^. while the isolated F2-F3 subdomains exhibit significantly higher binding affinity with integrin than the full size talin head domain, F2-F3 possesses limited activity in mediating integrin activation^15,16^. This underscores the crucial role of the structural integrity of talin head^16,24^. Interestingly, despite the conservation of the talin-binding motif in integrin-β, the structural basis for their widely varying affinities with talin remain elusive. It is also unclear how the FERM-folded talin head distinguishes conserved talin-binding motif in various integrin species to precisely orchestrate the association and signaling activity for diverse integrin functions.

To investigate the molecular basis underlying the activation of various integrin species by talin, we determined the crystal structure of talin head in complex with a peptide derived from the cytoplasmic tail of integrin β2. The structure unveils a similar FERM-domain conformation of talin head with the β2 peptide bound to the talin-binding groove with distinct side chain interactions. Moreover, in contrast to the β3-bound talin head, β2-bound talin head undergoes a significant “rocking” motion, reminiscent of the domain dynamics described in the “seesaw” model^15^. Structural and affinity measurement indicate that talin head in the β2-bound configuration binds to integrin with a lower affinity than in the β3-bound configuration. Notably, while keeping the interacting residues between talin and integrin intact, mutations restraining the talin head from the β2-bound configuration enhance talin:β2 association. Structural studies indicates that the mutant binds to β2 in the same configuration seen in the β3-bound talin. These observations lead to a hypothesis that stability of the C-terminal α-helix (C-α) in talin head contributes to its enhanced association with integrin. To explore this further, we engineered mutations in talin to stabilize the C-α helix. Talin bearing the mutations exhibits stronger association with integrin and enhanced activity in inducing integrin activation. Thus, the two configurations of the talin head domain exhibit distinct activities in mediating integrin activation, representing different preferences for recognizing β2 and β3 integrins. Our results, for the first time, offer a structural rationale for species-specific integrin activation by talin, providing crucial insight for developing targeted therapies towards various integrin species with greater specificity.

## Results

### Crystal structure of talin head in complex with β2 reveals a FERM conformation

Integrins engage with various cellular adapter proteins during activation and signaling. The majority of the integrin species possess both talin-binding and kindlin-binding sites within their cytoplasmic tail of the β subunit. Sequence alignment of this segment across various integrin β subunits reveals that both talin– and kindlin-binding sites harbor an NpxY/F motif, commonly found in transmembrane receptors to facilitate signaling transduction through protein-protein interactions (**Fig.1A**). Previous structural studies of β1 and β3 integrins have demonstrated that the talin:integrin-β association is mainly mediated by an 11-residue region centered around an NPxY/F motif. Interestingly, the corresponding region in β2 integrin is rather distinct, with a phenylalanine residue replacing the tyrosine residue in the NPxY/F motif, and one fewer residue compared with β1 and β3 (**Fig. 1A**). To elucidate the binding specificity of the β2 region and talin head, we generated a fusion protein comprising the talin-binding NPxY/F region from β2 and a full-length talin head lacking the F1-loop, and determined the crystal structures at 1.9 Å (**Table 1**).

**Table 1.** Data collection and refinement statistics.

|  | **β2:THD**<br>**(PDB:8FSE)** | **β2:THD(K306**<br>**Q)**<br>**(PDB:8FTB)** | **β2:THD(QAA)**<br>**(PDB:8T0D)** | **β2:THD(D397R)**<br>**(PDB: XXXX)** | **β3:THD(D397R)**<br>**(PDB: 9C1T)** |
| --- | --- | --- | --- | --- | --- |
| **Wavelength (Å)** | 0.92010 | 0.92011 | 0.92010 | 0.97934 | 0.97931 |
| **Resolution range**<br>**(Å)** | 43.85 - 1.90<br>(1.97 - 1.90) | 29.09 - 1.97<br>(2.04 - 1.97) | 29.04 – 2.77<br>(2.87 – 2.77) | 33.44 - 1.741<br>(1.804 - 1.741) | 29.78 - 2.30<br>(2.39 - 2.30) |
| **Space group** | P 1 21 1 | P 21 2 21 | P 21 2 21 | P 21 21 2 | P 21 21 21 |
| **Unit cell** | 66.86, 110.36,<br>68.37,<br>90°, 90.044°,<br>90° | 67.01, 68.39,<br>110.62,<br>90°, 90°, 90° | 67.11, 67.09,<br>115.84,<br>90°, 90°, 90° | 110.40, 66.89,<br>68.23,<br>90°, 90°, 90° | 66.43, 67.21,<br>115.57,<br>90°, 90°, 90° |
| **Unique reflections** | 77429 (7398) | 36623 (3575) | 13831 (1353) | 52004 (4994) | 23150 (1958) |
| **Multiplicity** | 3.2 (2.4) | 6.9 (6.9) | 13.0 (12.4) | 2.0 (2.0) | 2.0 (2.0) |
| **Completeness (%)** | 99.26 (95.68) | 99.78 (98.65) | 99.8 (99.9) | 99.09 (96.19) | 98.45 (85.27) |
| **Mean I/sigma(I)** | 8.4 (1.2) | 8.7 (1.9) | 17.0 (3.6) | 5.49 (0.96) | 12.68 (2.31) |
| **Wilson B-factor** | 19.09 | 23.66 | 55.95 | 19.21 | 47.03 |
| **R-merge (%)** | 13.3 (43.2) | 15.0 (81.9) | 15.6 (81.2) | 8.7 (69.2) | 3.1 (30.3) |
| **CC1/2** | 0.963 (0.666) | 0.994 (0.78) | 0.998 (0.876) | 0.993 (0.465) | 0.999 (0.812) |
| **Reflections used**<br>**in refinement** | 77429 (7397) | 36623 (3574) | 13829 (1353) | 51998 (4994) | 23145 (1956) |
| **R-work** | 0.1791 (0.2637) | 0.1833 (0.2195) | 0.1856 (0.2438) | 0.1981 (0.3136) | 0.2033 (0.2794) |
| **R-free (5%)** | 0.2199 (0.3251) | 0.2107 (0.2713) | 0.2416 (0.3299) | 0.2243 (0.3788) | 0.2669 (0.3424) |
| **non-hydrogen**<br>**atoms** | 7063 | 3539 | 3040 | 3724 | 3132 |
| **Protein** | 6079 | 3050 | 2995 | 3066 | 3046 |
| **Water** | 984 | 489 | 45 | 658 | 81 |
| **R.M.S Bonds (Å)** | 0.007 | 0.008 | 0.009 | 0.007 | 0.009 |
| **R.M.S Angles** | 0.81° | 0.98° | 0.97° | 1.00° | 0.96° |
| **Ramachandran**<br>**favored (%)** | 98.66 | 97.86 | 97.34 | 98.67 | 97.06 |
| **Ramachandran**<br>**outliers (%)** | 0.00 | 0.00 | 0.00 | 0.00 | 0.00 |

Statistics for the highest-resolution shell are shown in parentheses.

**Table 2.**
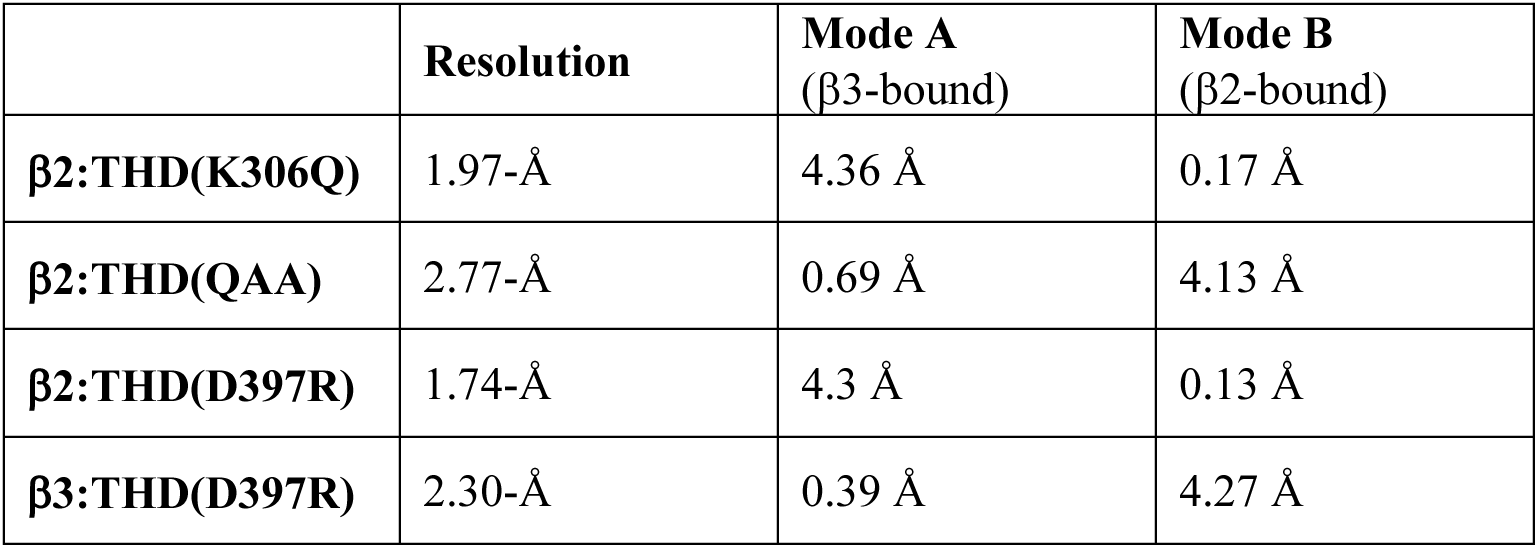
Comparison of the talin mutant structures with the two configuration modes. RMSD, listed below, were calculated by aligning all residues of the two structures.

The structure reveals two β2-THD molecules in an asymmetric unit. The two molecules are virtually identical with an overall root-mean-square deviation (RMSD) of 0.06-Å. The talin head domain folds into a FERM conformation, with the integrin-binding F3 subdomain positioning on top of the F1 and F2 subdomains, similar to that of the β3-bound talin head domain^24^. In both molecules, the fused β2 region engages with the talin head domain at the β5:C-α groove from a neighboring molecule (**Fig. 1B**). All ten residues of the β2 region are well resolved in the electron density (**Fig. 1C**). The interaction of the β2 and talin is largely attributed to hydrophobic interactions involving the Pro752, Leu753, and Phe754 residues of the NPxY/F motif with talin-F3 residues in β5 (Lys357), β6-β7 loop (Tyr373) and C-α (Ile396 and Leu400) (**Fig. 1D**). Additionally, Van-der-Waals and ionic interactions are observed via Arg358, Trp359, Asp372, and backbone of β4-β5 loop from the F3 subdomain (**Fig. 1D**). The number of residues in β2 between the NPxY/F motif and the juxtamembrane (JM) is one fewer than in other β subunits such as β1 and β3 (**Fig. 1A**). To evaluate the structural impact of this shortened connection, we analyzed the bound β2 with a β1-D structure that contains the JM region (PDB 3g9w). The structural comparison reveals that while the NPxY/F motifs of β2 and β1-D overlap, the three residues (Asn748, Asn749, and Asp750) preceding the motif in β2 extend to align Asn748 and Asn749 with Asp776 and Thr777 of β1-D. Consequently, the backbone stretching of the shortened connection residues would allow proper interaction of the JM region of β2 with talin (**Fig. 1E**).

### Subdomain tweaking in the talin head FERM domain leads to weakened integrin:talin interaction

Integrin species often exhibit tissue-specific expression patterns and vary in their affinity for talin and other adaptor proteins. We investigated the affinities of β2 integrin with talin using a fluorescence polarization (FP) assay with a FITC-labeled peptide representing both the JM and the NPxY/F regions, along with a purified talin head protein, THD405d (residues 1-405, lacking F1-loop). The β2 peptide exhibits a moderate affinity of 31.9 μM, which is significantly lower than the affinity of the β3 peptide, which was measured at 4.4 μM (**Fig. 2A**). The sequence variations between the β2 and β3 subunits undoubtedly lead to be observed differences in affinity. Additionally, we investigated how talin adapts to these differences in the β subunit, exploring its capacity to recognize various integrin species to ensure proper function.

**Figure 2.**
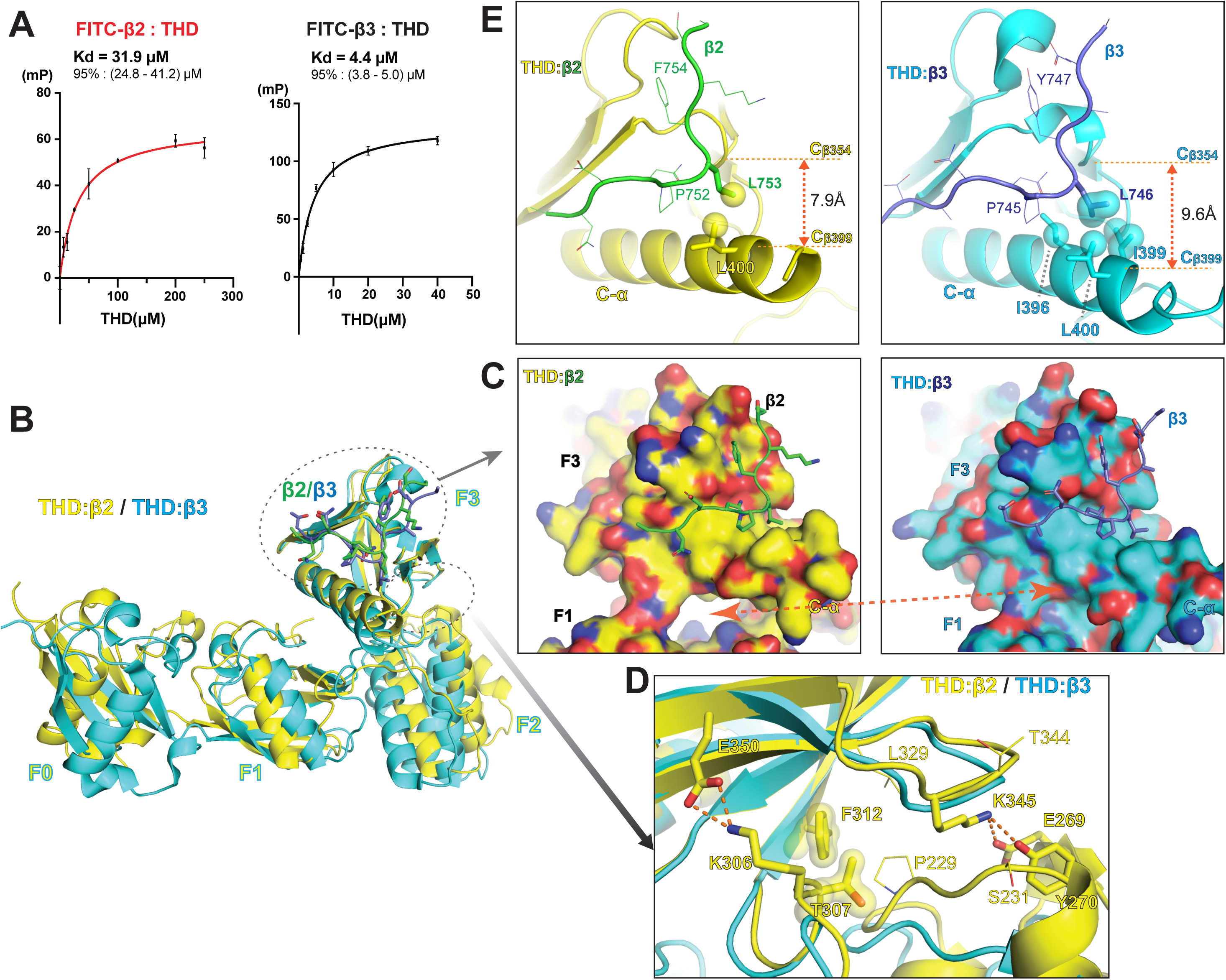
Comparison of the talin head binding to β2 and β3 integrins. **A**. Binding affinities of THD with β2 and β3 integrins were measured by FP assays. Calculated affinities and the 95% confidence interval are shown above the fitting curves. **B**. Superposition of the β2-bound THD and β3-bound THD. Alignment was performed using the F3 subdomain. **C**. Surface representation of the β2-bound THD (left) and the β3-bound THD (right). The F1-F3 cavity in the β2-bound THD, but absent in the β3-bound THD, is indicated by the double-headed arrow in red. **D**. Comparison of the F2-F3 interface in the β2-bound THD and the β3-bound THD reveals a tightly interacting F2-F3 interface in the β2-bound THD. Ionic interactions are indicated by dotted line. Other interactions are indicated by surface representation. **E**. Comparison of the hydrophobic interactions in β2:THD and β3:THD mediated by the Leucine residues (L753 in β2 and L746 in β3) in the NPxY/F (x=Leu) site. Note the integrin binding pocket is narrowed in β2:THD (left), indicated by the distance of Cβ354 and Cβ399, diminishing the hydrophobic interactions of the Leucine with I396 and I399 seen the in β3:THD structure (right).

Despite the overall similarity of the THD in the β2-bound form and the β3-bound form, the orientation of the subdomains changes noticeably. Superposition of the integrin-binding F3 subdomains of β2-bound THD and β3-bound THD reveals a significant movement in the F1 and F2 subdomains (**Fig. 2B**). In the β2-bound THD, the F1 subdomain shifts away from F3, creating a cavity between the F1 and F3 subdomains, while in the β3-bound THD, the two subdomains are in close contact (**Fig. 2C**).

Additionally, the F2 subdomain moves closer to F3 when bound to β2, leading to several new side chain interactions, including Lys306:Glu350, Thr307:Phe312, Glu269/Tyr270:Lys345 at the extended F2-F3 interface (**Fig. 2D**).

The tweaking of the THD subdomains induces in a shift in the C-α of the F3 subdomain towards the β5 strand, leading to a notable reduction in the size of C-α:β6 groove. Specifically, the distance from the Cβ atom of Ile399 in C-α to Thr354 Cβ in the β4-β5 loop reduces from 9.6-Å to 7.9-Å, resulting in a 5.7° narrowing of the C-α:β6 groove. This change hinders the interaction with the Leucine residue in the NPxY/F motif (where x=Leu in β2 and β3 integrins). In the β3:THD structure, Leu746 of β3 forms a robust hydrophobic cluster with Ile397, Ile399, and Leu400 of THD, whereas in the β2:THD structure, the corresponding Leu753 for β2 is unable to penetrate the C-α:β5 groove deeply enough to engage the hydrophobic sides chains of Ile396 and Ile399, thus retaining only the interaction with Leu400 side chain (**Fig. 2E**). Talin residues interacting with Asn751, Pro752, and Phe754 of the NPxY/F motif in β2 remain unchanged as those interacting with the corresponding sites in β3. These findings suggest that the C-α shift induced by the subdomain tweaking in THD is the primary cause of the weakened affinity with β2. It appears that FERM-folded THD may accommodate different integrin species through subdomain tweaking. These structural adjustments illustrate how talin’s flexibility enables it to recognize and interact with integrins in a species-specific manner, thereby contributing to proper cellular functions.

### Mutations at the F2-F3 interface modify the talin:integrin binding mode

We refer to the configuration of β3-bound THD as “Mode A” and the configuration of β2-bound THD as “Mode B” (**Fig. 3A**). In the “Mode A” configuration, the F3 subdomain tilts to the F1 subdomain, allowing the F1 subdomain to engage and stabilize the C-α helix. Specifically, Ile398 in C-α is sandwiched by Phe197 and Tyr199 via hydrophobic interaction, and Lys401 also forms a salt bridge interaction with Asp125 in the F1 subdomain. These interactions are disrupted in the “Mode B” configuration, as the F3 subdomain shifts towards the F2 subdomain. This shift results in an extensive F2-F3 interface, and the C-α helix retracts from the F1 subdomain, narrowing the integrin binding groove in F3. The tweaking of the subdomains between these two modes is indicated by an RMSD of 4.42 Å.

**Figure 3.**
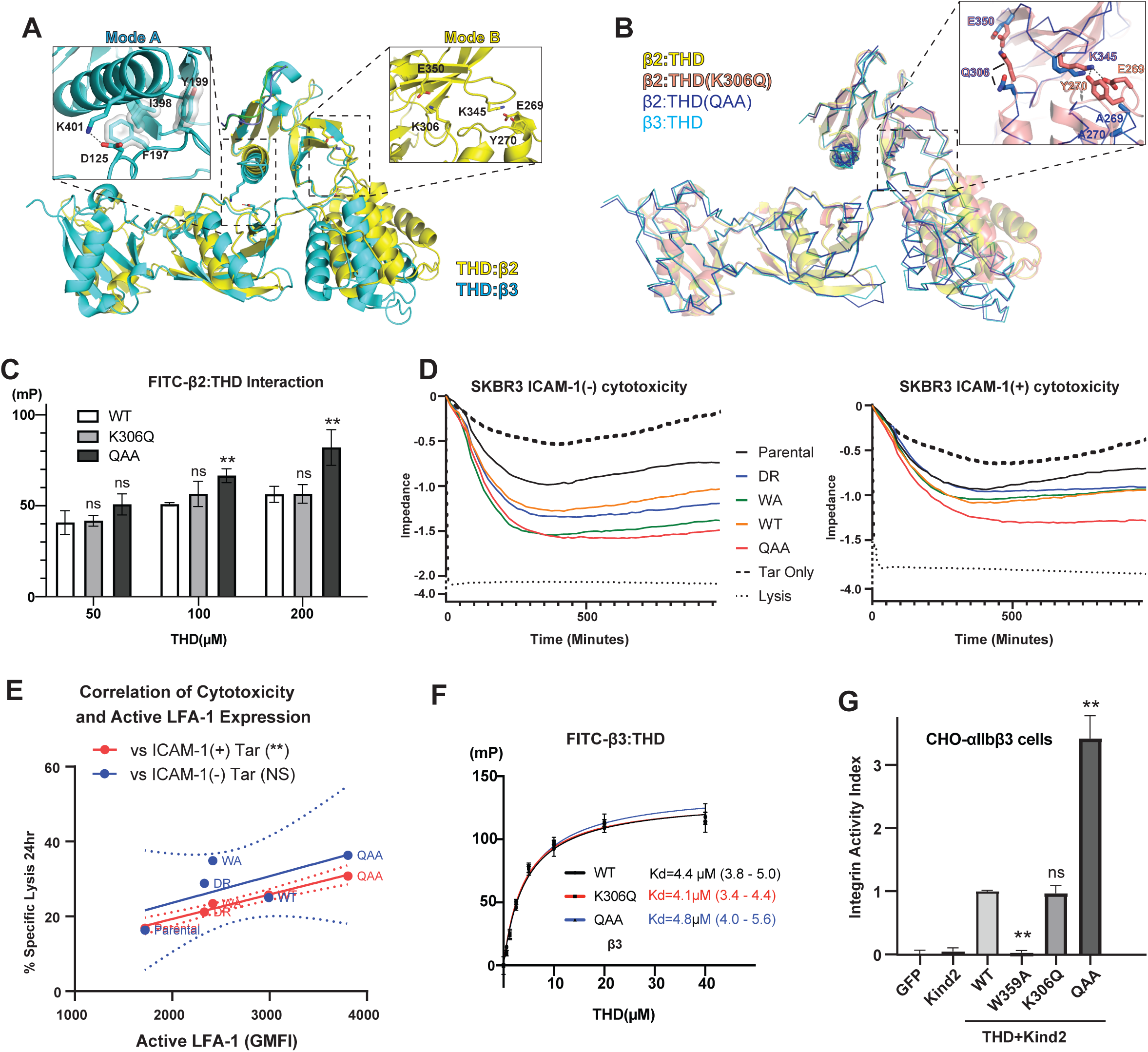
F2-F3 interface mutations alters talin:integrin-β binding mode and integrin activity. **A.** The β3:THD structure reveals a “Mode A” configuration of talin head (cyan), featuring a tight F1-F3 interface (left box), whereas the β2:THD structure reveals a “Mode B” configuration (yellow), featuring a tight F2-F3 interface (right box). **B**. Superposition of the crystal structure of β2:THD, β2:THD(K306Q), β2:THD(QAA), and β3:THD. β2:THD and β2:THD(K306Q), adopting the Mode B configuration, are shown in cartoon in yellow and salmon, respectively. β2:THD(QAA) and β3:THD, adopting the Mode A configuration, are shown in lines in blue and cyan, respectively. Superposition of the β2:THD(K306Q), β2:THD(QAA) at the F2-F3 interface, shown in the upper right box, indicates that the QAA triple mutation, but not the K306Q single mutation, disrupts the F2-F3 interface. **C**. THD:β2 interaction assessed by the FP assay. **D**. *Left:* cytotoxicity of NK-92 cells expressing various THD constructs against untreated SKBR3 cells (ICAM-1(-)), or *right:* against IFNγ-treated SKBR3 cells (ICAM-1(+)). **E.** Correlation of NK cell cytotoxicity and active LFA-1 level. **F.** THD:β3 interaction assessed by the FP assay. **G**. αIIbβ3 integrin activation induced by talin head is examined by the FACS assay. ns: Not significant. **: *p*<0.01

To assess the impact of the F2-F3 interaction on THD’s configuration, its association with integrin, and role in mediating integrin activation, we introduced mutations in THD, aiming to disrupt the F2-F3 interface. Particularly, a K306Q mutation is introduced to disrupt the Lys306:Glu350 salt bridge and a K306Q/E269A/Y270A (QAA) triple mutation is introduced to disrupt both Lys306:Glu350 and Glu269/Tyr270:Lys345 interactions. We determined the crystal structures of β2:THD(K306Q) and β2:THD(QAA) at 1.97-Å and 2.77-Å, respectively (**Fig. 3B**). The structure of THD:β2(K306Q) reveals a “Mode B” configuration of THD, nearly identical to that of the THD:β2, with an overall RMSD of 0.17 Å. This comparison suggests that the single mutation is insufficient to disrupt the F2-F3 interface when bound to β2 integrin. Strikingly, the structure of β2:THD(QAA) reveals a THD configuration of “Mode A”, similar to that of THD:β3, with an overall RMSD of 0.69 Å, but completely different from the THD:β2 (RMSD=4.13Å). As both Lys306:Glu350 and Glu269/Tyr270:Lys345 interactions are disrupted, the F2-F3 interface is opened as expected. We then compared the binding of β2 with the THD mutations using the FP assay. Although the affinity of β2 with each THD mutation could not be precisely measured due to limitations in both low affinity and protein solubility, the FP signal notably increases for the THD(QAA) mutation, whereas the FP signal change for THD(K306Q) is negligible (**Fig. 3C**). These observations indicate that disruption of the F2:F3 interface shifts talin head to the “Mode A” configuration, which affords a stronger association with the integrin β2 than the “Mode B” configuration.

### Shifting talin from “Mode B” to “Mode A” enhances LFA-1-mediated cytotoxicity by natural killer cells and αIIbβ3 integrin activity

β2 integrins are highly expressed in leukocytes. In NK (Natural Killer) cells, activated αLβ2 integrin (also known as LFA-1, or Lymphocyte function-associated antigen-1) mediates firm intercellular adhesion and signaling by clustering upon interaction with ICAM-1 (Intercellular Adhesion Molecule 1) on target cells at the lytic synapse ^25^. To investigate the impact of different binding modes on β2 integrin function, we assessed the cytotoxicity of NK cells against SKBR3, a breast cancer cell line, that lacks appreciable expression of ICAM-1 in the resting state. We induced ICAM-1 expression in SKBR3 through 48 hours of IFNγ treatment^26^ (**Fig. S1A**). We observed a dose dependent increase of ICAM-1 expression from 1-1000U/mL IFNγ. The various talin head constructs were then transduced by retrovirus into the human NK-92 cell line, which is functionally comparable to primary NK cells^27,28^. The cytotoxicity of NK cell lines against both untreated ICAM-1(−) and IFNγ-treated ICAM-1(+)SKBR3 target cells were monitored over time using xCELLigence technology (**Fig. 3D**). NK-92 cells were added to wells containing the adherent target cells and cytotoxicity was measured as loss of impedance signal on the culture well surface. Percent lysis of target cells by NK cells was determined by dividing the change in impedance at 24 hours by the total lysis control (**Figs. S1B and S1C**). Target cell lysis was increased to varying degrees by NK-92 cells transduced with various talin head constructs, compared to parental (untransduced) NK-92 cells. We then measured the total and active LFA-1 expression on the surface of these cell lines through FACS and found that the geometric mean fluorescent intensity (GMFI) of active LFA-1 expression was considerably different between NK-92 cells expressing various talin constructs (**Fig. S1D, S1E**). We then plotted the GMFIs of active LFA-1 expression in each of the NK-92 cell lines against their killing response and performed linear regression analysis (**Fig. 3E**). We observed a very strong positive correlation between active LFA-1 expression and the degree of killing against IFNγ-treated ICAM-1(+) SKBR3 cells (p=0.002, R^2=0.98) but not against untreated ICAM-1(^) SKBR3 cells (p=0.19, R^2=0.48). Notably, NK cells expressing the QAA mutation exhibited the highest expression of active LFA-1 and a significantly enhanced cytotoxicity. This finding aligns with the increased β2 affinity of THD(QAA) and supports that the mode A configuration of talin represents a stronger integrin binding state, leading to a higher β2 integrin activity.

We also examined the binding affinities of THD mutations with the integrin β3 peptide. Wild type THD, THD(K306Q) and THD(QAA) exhibit similar affinities, supporting the notion that these mutations would not affect the “Mode A” configuration (**Fig. 3F**). We then assessed the impact of the mutations in talin’s function in mediating β3 integrin activation in a fluorescence activated cell sorting (FACS) assay. CHO (Chinese Hamster Ovary) cells expressing αIIbβ3 integrin were co-transfected with GFP-labeled THD and kindlin-2, then examined for integrin activity in response to THD expression. Interestingly, although the THD(K306Q) mutation induces integrin activation to a similar level as the wild type THD, THD(QAA) induces αIIbβ3 integrin activation to a higher level **(Fig. 3G, Fig. S2)**. Thus, the K306Q/E269A/Y270A triple mutation disrupts the F2-F3 interface, facilitating the “Mode A” configuration of THD. Moreover, although the QAA mutation that disrupts the F2-F3 interface does not affect its β3-bound configuration (Mode A) and β3 affinity, it restrains the dynamics of the FERM domain and locks talin head in the “Mode A” configuration. These findings suggest that the “Mode A” configuration in which the C-α helix is stabilized by F1, represents a state of talin with both high affinity and high activity. Since integrin β3 activation requires simultaneous association with both talin and kindlin, we also examined the talin:β3 interaction in the presence of kindlin-2. While the presence of kindlin-2 does not change the association of β3 with wild type talin, its association with talin THD(QAA) is significantly enhanced (**Fig. S3**), consistent with the elevated integrin activation induced by THD(QAA). Thus, the equilibrium shift from “Mode B” to “Mode A” of THD may lead to increased integrin activity without altering the apparent affinity with integrin.

### Stabilization of talin head C-α enhances integrin association and talin-induced integrin activation

To further investigate the role of C-α in integrin association, we proposed that the stability of C-α contributes to a stronger association with integrin. To test this hypothesis, we introduced mutations with the goal of stabilizing the C-α helix. We mutated Asp397 to a positively charged side chain to promote intra-helix interaction with other residues within C-α or inter-subdomain interactions with residues from F1. If successful, these additional interactions would enhance the stability of the C-α helix. To validate the additional interaction, we determined the crystal structure of β3:THD(D397R). The structure reveals a “Mode A” THD configuration with an RMSD of 0.39 Å when compared with the wild-type THD when bound to β3 (PDB code: 6vgu) (**Fig. 4A**). Remarkably, the mutated residue Arg397 forms an intra-helix hydrogen bond with Gln390, forming a side-chain “staple” that stabilizes the C-α helix (**Fig. 4A**). We then assessed the association of the mutant THD with integrins. Both THD(D397R) or THD(D397K) demonstrated a two-fold increase in affinity for β3 integrin compared to the wild-type THD, as indicated by the FP assay (**Fig. 4B)**. In the truncated talin head domain (talin-400), which has impaired activity in its linear configuration, the talin-400(D397K) and talin-400(D397R) mutants also showed enhanced association with β3 integrin (**Fig. S4A**). Further cellular function analyses indicated that co-expression of the THD(D397R) mutant with Kindlin-2 in CHO cells resulted in greater activation of integrin αIIbβ3 compared to the wild-type THD (**Fig. 4C, Fig. S2**). Nevertheless, these mutants did not restore activity to the impaired talin-400 the activity of the impaired talin head (**Fig. S4B, C**), indicating that the mutations do not facilitate the transition from a linear to a FERM-folded configuration necessary for integrin activation by positioning the F1 loop to engage and separate the integrin α and β subunit^29^.

**Figure 4.**
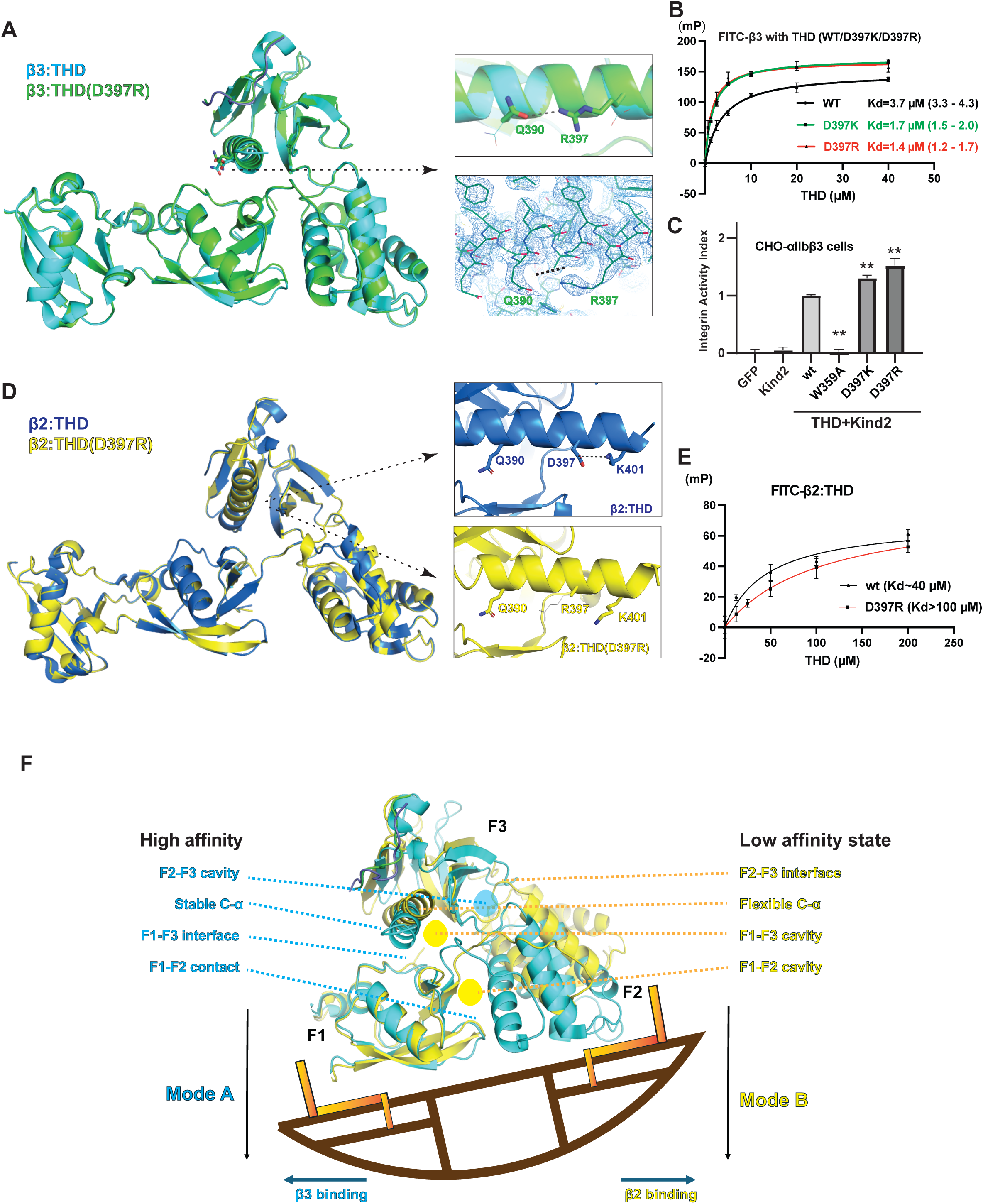
Stabilization of the C-α helix in talin F3 contributes to the THD:integrin-β interaction. **A**. Crystal structure of β3:THD(D397R). *Left*: superposition of β3:THD (cyan) and β3:THD(D397R) (green) structures; *Upper right*: Residues Q390 and R397 in the β3:THD(D397R) structure, shown in stick representation in green, form a hydrogen bond interaction. Corresponding residues Q390 and D397 in the wild type β3:THD structure are shown in line representation in cyan. *Lower right*: electron density of the C-α helix of the β3:THD(D397R) structure contour at α=1.0. **B**. THD:β3 interaction assessed by the FP assay. Both THD(D397K) and THD(D397R) variants enhance the THD:β3 interaction. *Kd* for β3 with each protein was calculated, with the 95% confidence interval provided in parentheses. **C**. CHO-αIIbβ3 cells expressing either kindlin2 alone (Kind2) or together with talin head constructs (THD+Kind) were examined for αIibβ3 activity by FACS. **D**. Crystal structure of β2:THD(D397R). *Left*: superposition of β2:THD (blue) and β2:THD(D397R) (yellow) structures; *Upper right*: a salt bridge interaction from by residues D397 and K401in the β2:THD structure. *Lower right*: The side chain of R397 in the β2:THD(D397R) structure beyond C-alpha (stick representation) is disordered (shown in line representation), mediating no side-chain interaction. **E.** THD:β2 interaction assessed by the FP assay. Affinity of THD(D397R) with β2 is significantly reduced (*Kd* > 100 μM). **F**. A rocking seesaw model of the species-specific recognition of integrin by talin. Only the F1, F2, and F3 subdomains of talin head are shown. In Mode A (cyan), the F1:F3 interface stabilizes the C-α, and F1 contacts F2, leading to a tighter β3:talin interaction. The Mode B (yellow), F3 rocks to the F2 subdomain side, forming the F2:F3 interface and generating F1:F3 and F1:F2 cavities, resulting in a flexible C-α and a weaker β2:talin interaction. **: *p*<0.01

We also determined the crystal structure of β2:THD(D397R). The structure reveals that talin remains in the “mode B” configuration and the disordered Arg397 side chain forms no additional interaction (**Fig. 4D**). Interestingly, in the wild type β2:THD complex, the Asp397 residue forms a salt bridge with Lys401 (**Fig. 4D**, *upper right*), and the D397R mutation diminishes the Asp397:Lys401 interaction in the β2-bound THD instead of generating a new intra-helix interaction as seen in the β3-bound structure (**Fig. 4D**, *lower right*). FP assay indicates that the D397R mutation reduces the β2 affinity to talin (**Fig. 4E**), consistent with the proposed model as D397R destabilizes the C-α helix in the “mode B” configuration. Consequently, in the LFA-1 mediated cytotoxicity assay, talin(D397R)-expressing NK cells exhibit impaired cytotoxicity to ICAM-1^high^ SKBR3 cells due to the impaired integrin function (**Fig. 3E**). These results further support the hypothesis that stabilizing the C-α helix enhances the interaction between talin and integrins.

## Discussion

Understanding how talin associates with different types of integrins is crucial for unraveling the complex processes of tissue-specific cell adhesion and migration. It also helps identify new targets for therapeutic intervention and development of more specific treatments that reduce side effects caused by cross-reaction with other integrins. β3 integrin, when partnered with different α subunits, plays diverse roles in various cell types, including platelet aggregation (αIIbβ3), bone resorption (αVβ3), and contributes to pathological conditions such as cardiovascular diseases and cancer metastasis. In contrast, β2 integrins are predominantly expressed on leukocytes and platelets and play crucial roles in immune responses by pairing with an exclusive set of integrin α subunits. αLβ2 (LFA-1) is widely expressed on lymphocytes and plays essential roles for lymphocyte migration and T-cell activation. αMβ2 (Mac-1) is abundantly expressed in neutrophils and monocytes and mediates phagocytosis and the inflammatory response ^19^.

αXβ2 (CD11c/CD18) is mainly found on dendritic cells and contributes to antigen presentation and cell adhesion ^20^. αDβ2 (CD11d/CD18) is strongly expressed on some specific macrophages and may play a role in phagocytosis and atherosclerotic process ^30^. Together, these beta2 integrins mediate various critical immune processes, including leukocyte trafficking, phagocytosis, and immune cell activation.

Our findings reveal different binding modes and affinities of talin when associates with β2 and β3 integrins. In hematocytes, β3 integrin is abundant in platelets, β2 integrin is prevalent in lymphocytes. Platelets respond rapidly to injury by forming blood clots, whereas lymphocytes take longer to initiate precise and necessary immune responses. Perhaps the high-affinity talin:β3 binding mode and the low-affinity talin:β2 binding mode may reflect the functional requirements of these cells: platelets prioritize rapid activation and adhesion, whereas lymphocytes focus on specificity and accuracy in immune responses.

While the crystal structure of β2-bound THD reveals a conventional FERM-fold, it exhibits unique conformational characteristics in contrast to the β3-bound THD. With talin’s capacity to recognize integrins with diverse sequences and with post-translation modifications, the structural features of its two binding modes will offer new insight into talin’s specificity across different integrin species. Activated talin translocates to the plasma membrane with the assistance of Rap1, RIAM, or PIPs, allowing it to engage and capture the membrane-anchored integrin βsubunits with a “*K_on_*” rate. The “*K_on_*” rate is primarily determined by the abundance of integrins, the surface complementarity between the integrin β and the integrin-binding groove of talin, and the accessibility of the integrin-binding groove. Once talin captures an integrin, the dissociation rate, or *K_off_*, is determined by the stability of the complex. In talin:integrin complex, the flexibility of the C-α helix of talin F3 subdomain is the key factor. In binding mode A, C-αis stabilized by the F1 subdomain, resulting in a lower *K_off_*, and subsequently, a higher affinity for integrin β3. In contrast, in binding mode B, the C-α helix is rather isolated, leading to a higher *K_off_* and a relative lower affinity for integrin β2. In the case of the THD(QAA) mutant, where binding mode A is prioritized, only the affinity of β2, but not β3, is enhanced. This observation is further supported by the finding that the THD(D397R) mutation, which enhances the stability of the C-α helix in the β3-bound state while diminishing stability in the β2-bound state, demonstrates inverse effects on affinities and functions of β2 and β3 integrins. Integrin activation requires simultaneous association with talin and kindlin. The dynamics of talin, kindlin and integrin-β interaction remains to be elucidated.

Recent studies suggest an equilibrium for talin and kindlin to engage integrin-β and form a tertiary complex that is stabilized by a direct interaction of talin-F^31^. We also observed that kindlin-2 significantly enhances the interaction of integrin β3 with THD(QAA) (**Fig. S3**). These observations suggest that the mode A configuration, favored by the THD(QAA) mutant, represents an optimal talin configuration for cooperative activation with kindlin, leading to a higher integrin activity.

Our model indicates that talin may interact with various integrin species in different configurations. The difference between the β3-binding mode (mode A) and the β2-binding mode (mode B) focuses mainly on the subdomain interfaces. Mode A features closed F1:F3 and F1:F2 interfaces and an F2-F3 cavity, whereas mode B contains a closed F2:F3 interface and cavities in F1-F2 and F1-F3. The lack of F1:F3 contact in mode B also results in a flexible C-α in F3. These structural distinctions may be utilized to target integrins for therapeutic intervention and to identify pathological biomarkers with greater species specificity. Antagonists of integrins have been developed to inhibit integrin function by blocking the extracellular ligand binding site. Several antagonists that target integrins with high expression profiles in hematocytes have been approved for medical uses to treat cardiovascular diseases and various types of autoimmune diseases ^32^. The limitations of these antagonists are reflected in severe adverse effects such as thrombocytopenia, hemorrhage, and the lethal Progressive multifocal leukoencephalopathy (PML) ^32,33^ caused by unexpected agonism and reactivation of the JC Virus ^11,12,34–36^. The different binding modes adapted by talin when binding to platelet-rich β3 integrin and leucocyte-rich β2 integrin reveal unique cavities between subdomains, such as the F1-F2 and F1-F3 cavities in the THD:β2 complex and F2-F3 cavities in the THD:β3 (**Fig. 4F**). Particularly, we show that LFA-1-mediated NK cell cytotoxicity can be enhanced by modifying talin configuration. Thus, by stabilizing talin in specific conformations using small molecules, broader and more precise immune responses may be achieved. This approach offers a structural basis for designing compounds that target distinct binding sites within each talin configuration, optimizing integrin specificity and affinity. Moreover, combining these small molecules with extracellular antagonists or intracellular peptidomimetics could lead to more precise targeting of integrin subtypes, potentially reducing adverse effects while improving therapeutic efficacy.

In summary, understanding how talin associates with different integrins is critical for unraveling the complex mechanisms of tissue-specific cell adhesion, migration, and immune responses. Our study reveals the crystal structure of the talin head bound to the β2-integrin tail, demonstrating a two-mode configuration with distinct inter-subdomain interactions and binding dynamics compared to the β3-integrin interaction. These structural differences help explain talin’s varying affinities for β2 and β3 integrins, which are predominantly involved in immune and platelet functions, respectively. Our findings provide a structural basis for the development of integrin species-specific therapeutic agents by targeting talin in an allosterically manner to modify its integrin preferences. The combination of these agents with agents directly targeting integrins could lead to more precise and effective treatments for various integrin-related conditions, while minimizing potential side effects caused by current treatments.

## Materials and methods

### Plasmid construction

Talin head was subcloned into a modified pET28a vector with a His6-tag. The cytoplasmic tails of different β integrins were subcloned into pGEX-5X-1 vector with a GST-tag. Talin head was subcloned into an EFEP-C3 vector for expression in CHO-A5 cells that express αIIbβ3 integrins stably. For crystallization, a truncated cytoplasmic tail of human integrin β2 (residues 748-757) was fused to the N-terminal of the talin head domain spreading from residue 1 to residue 430 with the deletion of loop (residue 139-168) in the F1 subdomain^15^. The fusion protein was inserted into a modified pET28a vector, with the His6-tag and linker removed by mutagenesis. Point mutations were generated with a site-directed mutagenesis according to the QuikChange Site-directed mutagenesis manual. All constructs were confirmed by automated DNA-sequencing.

### Protein purification

The expression and purification of the His6-tagged proteins and GST-tagged proteins were carried out as described previously^24,37^. Briefly, *E. coli* BL21(DE3) was transformed with modified pET28a-THD constructs or pGEX-5X-1-integrin-β-tail constructs. Bacteria were cultured in LB medium containing 50 µg/ml kanamycin or 80 µg/ml ampicillin in shaking flasks at 200 rpm at 37℃ until the OD600nm reached 0.7. To induce protein expression, 0.2 mM isopropyl-D-1-thiogalactopyranoside (IPTG) was added to the flasks. The flasks were then incubated at 200 rpm, at 16℃ for His6-tagged protein production, or for 3h at 37℃ for GST-tagged protein production. Bacteria were collected by centrifugation and were resuspended in 20 mM Tris, pH 7.5, 500 mM NaCl for His6-tagged proteins, or 20 mM Tris pH 7.5, 100 mM NaCl, 2 mM dithiothreitol (DTT) for GST-tagged proteins. Resuspended cells were homogenized with EmulsiFlex-C3, and the supernatant was collected as the cell extracts. The cell extracts were clarified and applied to HisTrap FF columns (GE LifeSciences) or GSTrap HP columns (GE LifeSciences) for purification using an ÄKTA Purifier system (GE Healthcare).

The expression of untagged proteins (β2-THD, β2-THD(K306Q), β2-THD(QAA), and β3-THD(D397R)) was carried out as described for His6-tagged protein expression. The cell pellet was collected by centrifugation and subsequently resuspended with 20mM 4-(2-hydroxyethyl)-1-piperazineethanesulfonic acid (HEPES), pH 7.0, 80 mM NaCl, 2 mM DTT. The cell extract generated by EmulsiFlex-C3 was loaded onto Hi Prep SP XL 16/10 (Cytiva) for purification using an ÄKTA Purifier system (Cytiva).

### Crystallization and structure determination

Purified untagged proteins were concentrated to 30-45 mg/ml in 20 mM HEPES, pH 7.0, 200 mM NaCl, and 2 mM DTT. Crystallization was performed using the hanging-drop vapor diffusion method at room temperature. The original β2-THD crystal was obtained by seeding microcrystals of β3-THD into the hanging drops in the wells containing a well-solution of 100 mM MES pH 6.5, 100 mM NaCl, 10% PEG4000^15^. Crystals of β2-THD(QAA) and β2-THD(K306Q) were obtained by micro-seeding with crystals of β2-THD. Crystals of β3-THD(D397R) were obtained by micro-seeding using crystals of β3-THD. Crystallization conditions for each protein were further optimized before harvested for data collection. The β2-THD430d crystals were harvested from a well-solution containing 0.1M 2-(N-morpholino) ethanesulfonic acid (MES), pH 6.5, 14% (w/v) polyethylene glycol (PEG) 4000 and 0.12M NaCl. Crystals of β2-THD(K306Q) and β2-THD(QAA) were harvested from a well solution containing 20% (w/v) polyethylene glycol (PEG) 3350 and 0.25 M Li3Cit. The β3-THD(D397R) crystal for data collection was harvested from a well solution containing 0.1M 2-(N-morpholino) ethanesulfonic acid (MES), pH 6.5, 10% (w/v) polyethylene glycol (PEG) 4000 and 0.1M NaCl. X-ray diffraction data were collected at the Brookhaven National Laboratory NSLS-II AMX and FMX beamlines. The crystal structures were determined by molecular replacement using β3-THD structure (PDB:6VGU) as a model. The structures were refined by COOT and Phenix. The final atomic coordinates and structure factors have been deposited to Protein Data Bank with the following accession numbers: β2-THD (8FSE), β2-THD(QAA) (8T0D), β2-THD(K306Q) (8FTB), and β3-THD(D397R) (9C1T).

### GST pull-down

The pulldown was performed as described previously^15^. Purified His6-THD proteins and GST-β tail/ GST proteins were mixed in binding buffer (50 mM Tris, pH 7.5, 100 mM NaCl, 2 mM DTT) to a final volume of 100 μl at a concentration of 2.1uM and 3.6uM individually. The protein mixture was left on ice for 15 min then incubated with binding buffer-equilibrated glutathione agarose beads (Invitrogen, G2879) on a rotator for 1h at 4 ℃. After removing the supernatant, the beads were washed with binding buffer three times. The beads-binding proteins were eluted with 30ul elution buffer (50 mM Tris, pH 7.5, 100 mM NaCl, 2 mM DTT, 20 mM reduced glutathione). The samples were applied to SDS/PAGE and analyzed by Coomassie staining or Western Blotting. Anti-His (Sigma) was used for the detection of His6-tagged THD proteins. Anti-Kindlin-2 (CST) was used for the detection of the kindlin-2 protein. A FluorChem E System (Proteinsimple) with a charge-coupled device (CCD) camera was used to expose the Western blotting membrane.

### Fluorescent polarization

The Fluorescent polarization assay was carried out as previously described. (Gao structure). The talin head protein solutions (at a proper series concentration) or buffer (20 mM Tris pH 8.0, 100 mM NaCl, 2 mM DTT) were mixed with 20 nM FITC-β2 (Genemed Synthesis, Inc.) or FAM-β3 (Genemed Synthesis, Inc.) in 20 mM Tris pH 8.0, 100 mM NaCl, 2 mM DTT, 0.5% Tween-20. The mixture was incubated on ice for 3 min then 20 µl was aliquoted to a 384-well plate (Corning, 3573) for measurement with Perkin Elmer Envision Plate Reader. The excitation filter was FITC FP 480nm and emission filter was a pair of FITC FP P-pol 535nm. All the polarization signals were normalized by subtracting the background signal generated by the buffer and fit to a single-site (saturating) binding model using Prism 9. For FITC-β2 binding, the polarization signals generated by mixing 20nM FITC-β2 with 50µM, 100µM or 200µM talin head protein were normalized by subtracting the background signal and data represent the mean of triplicates with error bars representing ±SD.

### Integrin activation assays

The CHO-A5 cells that expressed αIIbβ3 integrins were transfected with EGFP-C3-THD constructs and incubated at 37℃ for 36H. The cells were detached with an enzyme-free cell dissociation solution (Sigma, c5914-100ml) and washed with PBS. Then CHO-A5 cells were incubated with PAC-1 antibody (Invitrogen, MA5-28523, 1:50) in Tyrode’s buffer (136.9 mM NaCl, 10 mM HEPES, 5.5 mM Glucose, 11.9 mM NaHCO3, 2.7 mM KCl, 0.5 mM CaCl2, 1.5 mM MgCl2, 0.4 mM, NaH2PO4, pH 7.4) for 30 min at RT, followed by incubation with Alexa 647 goat anti-mouse IgM (Jackson Immuno Research Labs, NC0401238, 1:400) in Tyrode’s buffer for 30min on ice in the dark. After washing, the cells were resuspended with PBS and quantified with a Symphony A5 analyzer (660/20 filter, 640 nm laser) using 30,000 cells per measurement. All the data were processed with FlowJo v9.

### Cell line construction

To create NK-92 lines expressing talin head constructs, Talin-1 head domain (residues 1-430) and its various mutations (DR:D397R, WA:W359A, and QAA:K306Q/E269A/Y270A) were cloned into pBMN-IRES-EGFP vector (Garry Nolan, Stanford University). Retroviral particles were then created using amphotropic Phoenix packaging line (Garry Nolan, Stanford University) that were then transduced into NK-92 cells as previously described^38^. After six days, stable EGFP-positive cells were sorted from transduced line in the FCCC Cell Sorting Facility and cultured for downstream assays.

### Cell killing assay

NK-92 cells were passed every 3-4 days in NK cell growth media consisting of Alpha-MEM, 100U/mL recombinant human IL-2 (teceleukin, Hoffman-La Roche Inc., generously provided by the Biological Resources Branch of NCI-Frederick) and other supplements, as previously described. SKOV3 and SKBR3 cell lines were cultured in either RPMI-1640 (SKOV3) or McCoy’s (SKBR3) medium supplemented with 10% FBS. For staining of LFA-1, total relative LFA-1 levels in cell lines were determined using an HI111 anti-CD11a antibody (BD), while active LFA-1 was stained using an m24 anti-CD11a/CD18 antibody (BioLegend) as previously reported^39^, then measured with FACS. To assess elevation of ICAM-1 levels on target cell line, SKBR3 cells were transferred to fresh growth media supplemented with 1 to 1000 U/mL of IFN-γ for 48 hours prior to any functional assays^40^. ICAM-1 levels were stained using the HA58 anti-ICAM-1 antibody (Invitrogen) and analyzed by FACS. To assess cytotoxicity of adherent target cells over time, xCELLigence real-time cell analysis (RTCA; Agilent Technologies) was employed, as previously described^41^. 48 hours after 100U/mL IFN-γ or vehicle treatment, cells were washed twice, disassociated using an EDTA-containing wash buffer, resuspended in growth media, and plated onto E-16 plates (10,000 cells/well). Cytotoxicity assays were performed the following day at a 5:1 effector to target (E:T) ratio of NK-92 cells in a 50:50 mixture of NK cell growth media and target cell growth media to minimize NK cell-independent disassociation of SKBR3 cells from the wells. Figure assembly was done in Prism v10.3 (Graphpad). Flow cytometry data was collected on a FACSAria II in the FCCC Cell Sorting Facility and analyzed in Flowjo X (BD). Linear regression analysis was done using the SciPy Python library.

## Statistics

The crystallographic data were processed and refined with COOT, PHENIX, and REFMAC. The data of fluorescent polarization assays were normalized by subtracting the background signal generated by the buffer only. In integrin activation assay, the background signal of GFP-transfected cells was subtracted and all the data were normalized to the integrin activity level generated by GFP-THDwt transfection. Data represent the mean of triplicates with error bars representing ±SD. The statistical significances were determined using unpaired two-tailed Student’s T-test.

## Acknowledgments

We thank Dr. Bernhard Wehrle-Haller (University of Geneva) for the discussions, and Dr. Jun Qin (Cleveland Clinic) for the CHO-A5 cells. We thank the beamline staff of AMX and FMX at National Synchrotron Light Source-II, Brookhaven National Laboratory, 7B2 at MacCHESS, Cornell University, for technical support. This work was supported by an NIH Grant GM119560 (to J.W.), an ASH bridge grant (to J.W.), a Pennsylvania Department of Health Grant 4100085739 (to J.W.), and ACS RSG-15-167-01-DMC (to J.W.). T.G. was partially supported by the Elizabeth Knight Patterson Postdoctoral Fellowship. S.K. was partially supported by the Jeanne E. and Robert F. Ozols Undergraduate Summer Research Fellowship.

## Author Contributions

T.G. and S.K. carried out protein production and the FP assays. T.G. performed protein crystallization, data collection, biochemical experiments, and the cell-based functional experiments. T.G. and J.W. determined the structures. N.A.M., K.S.C., and J.W. designed the cytotoxicity experiments with N.A.M. executing the experiments. N.A.M., S.K., and K.S.C contributed to the manuscript preparation. T.G. and J.W. wrote the manuscript. J.W. supervised the project and was the principal manuscript author.

---

**Supplemental Figure 1.**
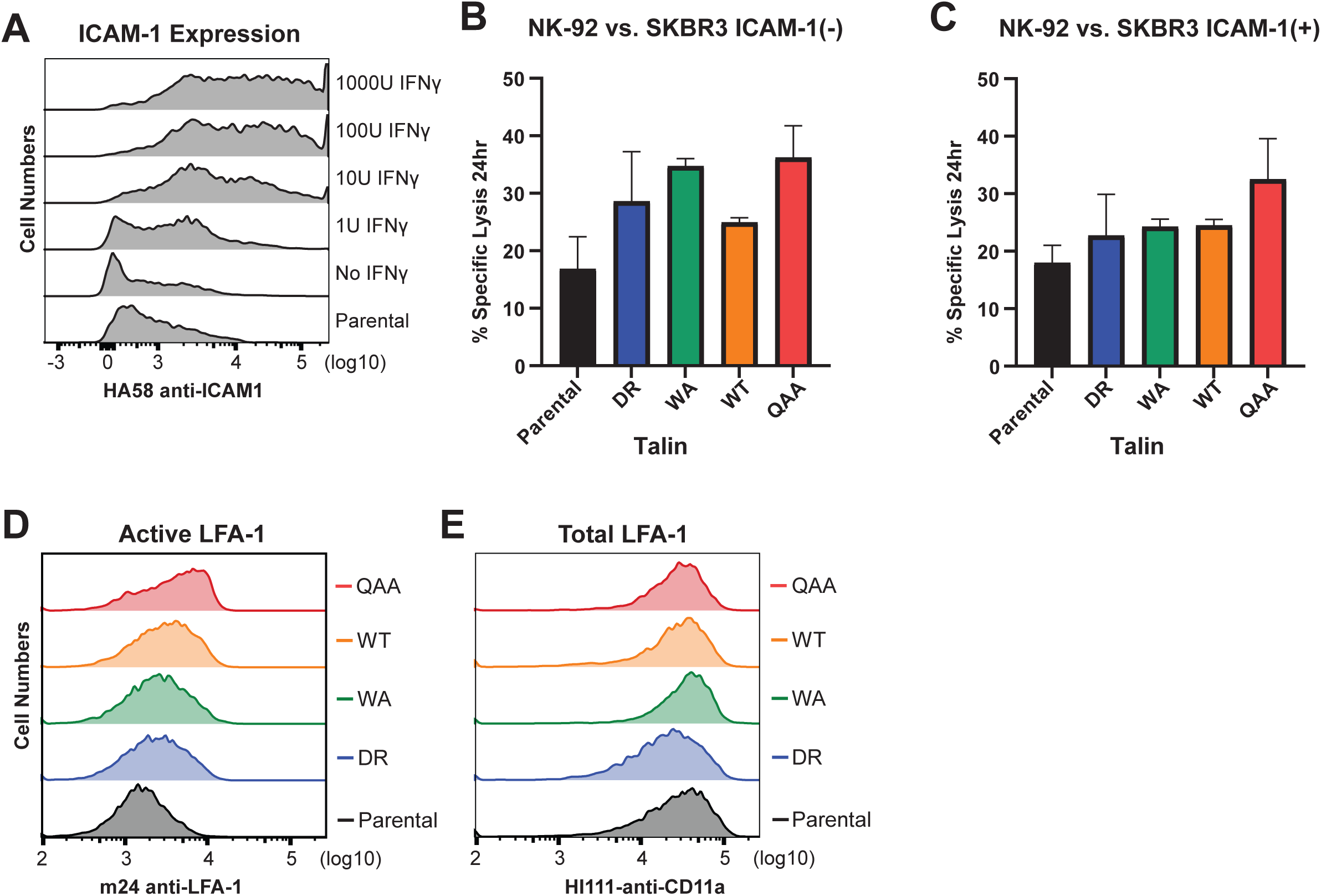
A. Expression of ICAM-1 in SKBR3 cells induced IFYɣ B. Present Iysis of the ICAM-1(-) SKBR3 cells by NK cells C. Present Iysis of the ICAM-1(+) SKBR3 cells by NK cells D. Active LFA-1 level in NK –92 cells expressing various talin constructs E. Total LFA-1 level in NK –92 cells expressing various talin constructs

**Supplemental Figure 2.**
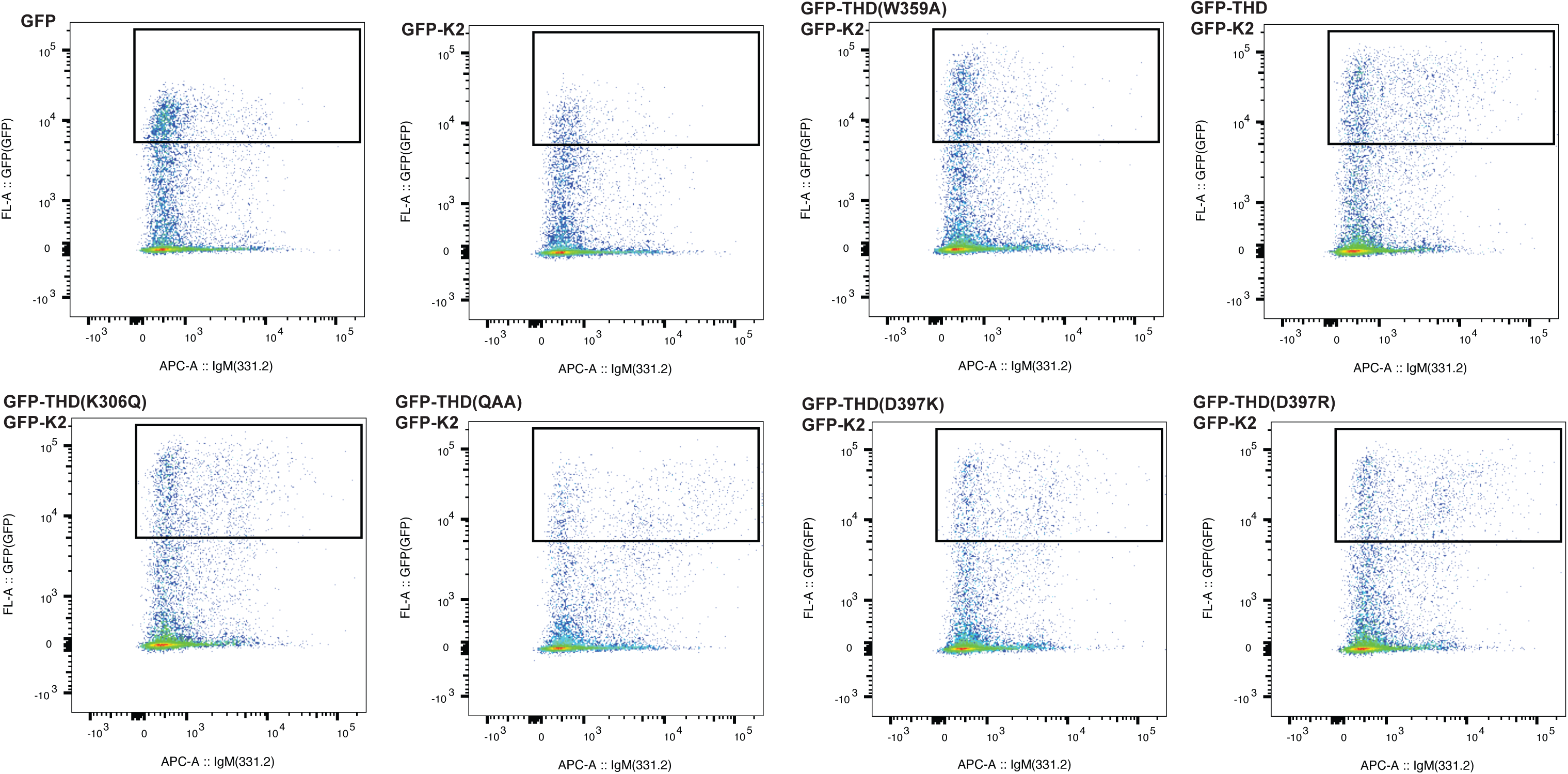
Fluorescence activated cell sorting (FACS) assay for CHO-aIIbf33 cells. Cells were transfected with the indicated plasmids. Cells with GFP fluorescence intensity greater than 10A3.5 were selected for PAC-I antibody analysis.

**Supplemental Figure 3.**
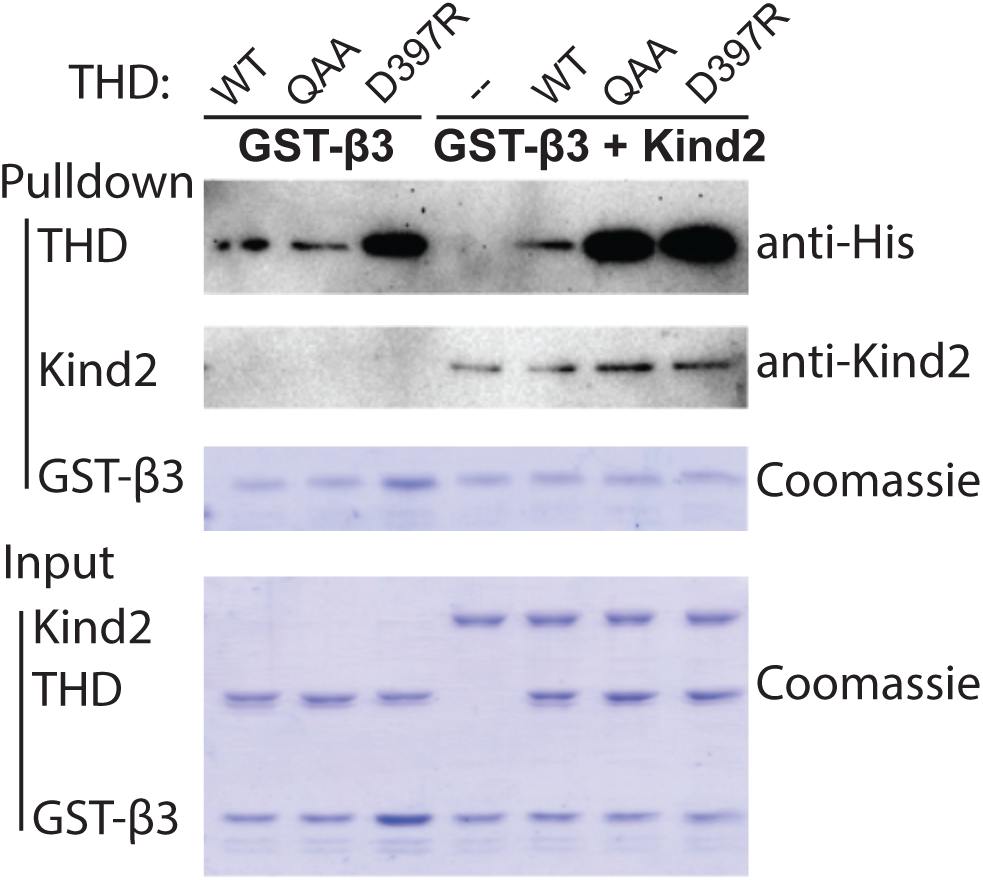
Pulldown of talin head domain and kindlin-2 by GST-f33 (residue 742-788).

**Supplemental Figure 4.**
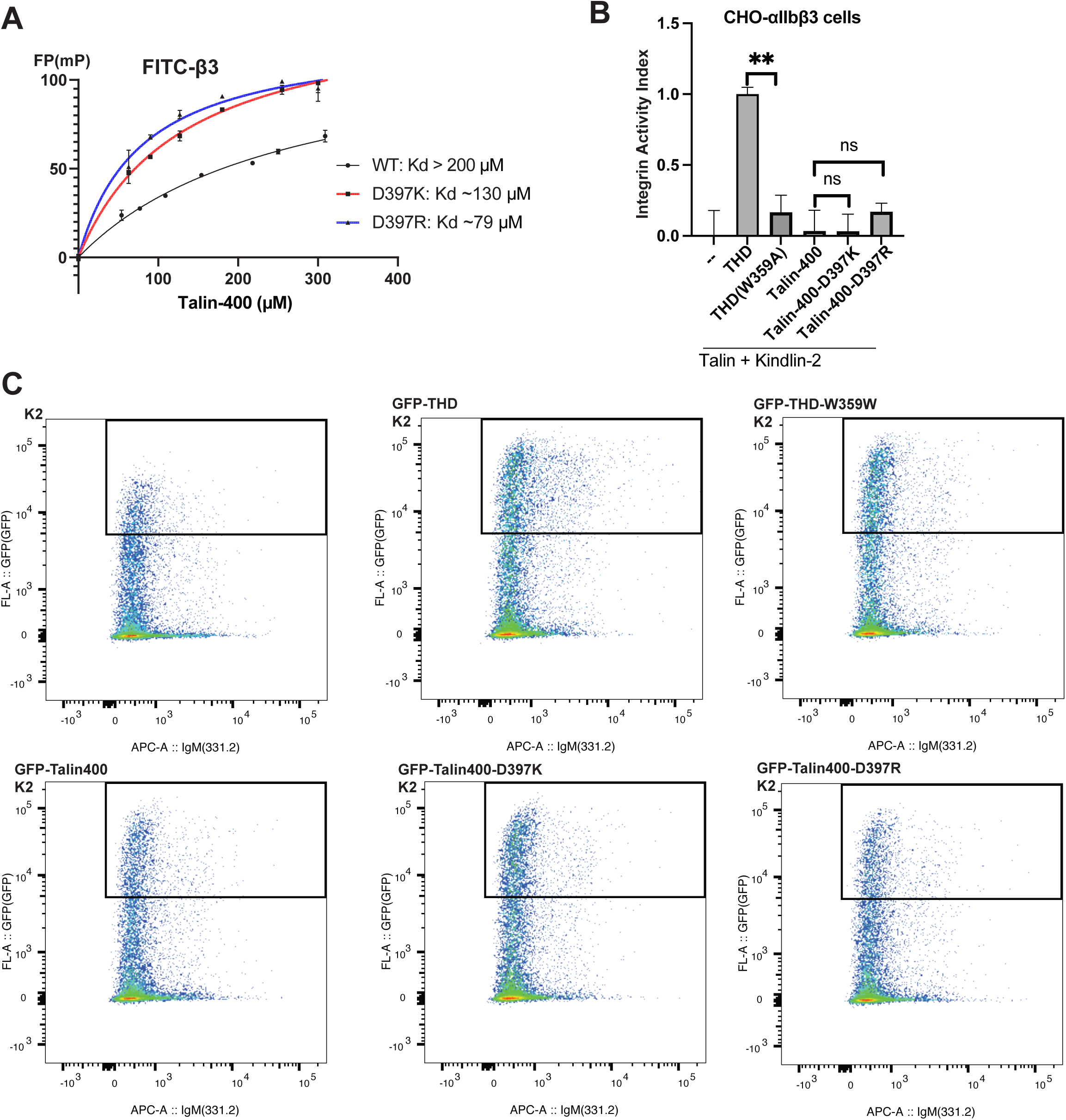
A. Affinities of Talin-400 constructs with integrin-f33 measured by FP. B. Integrin-f33 activation byTalin-400 proteins measures by FACS. C. Fluorescence activated cell sorting (FACS) assay for CHO-aIIbf33 cells. Cells were transfected with the indicated plasmids. Cells with GFP fluorescence intensity greater than 10A3.5 were selected for PAC-I antibody analysis.

